# Alcohol and Cannabinoid Binges and Daily Exposure to Nicotine in Adolescent/Young Adult Rats induce Sex-Dependent Long-Term Learning and Motivation Alterations

**DOI:** 10.1101/2022.12.20.521255

**Authors:** Norbert Abela, Katie Haywood, Giuseppe Di Giovanni

**Affiliations:** Laboratory of Neurophysiology, Department of Physiology and Biochemistry, Faculty of Medicine and Surgery, University of Malta - Msida, Malta; School of Biosciences, Neuroscience Division, Cardiff University, Cardiff, UK

**Keywords:** Marijuana, smoking, adolescent drug abuse, learning, cannabinoids, nicotine

## Abstract

Adolescence is a critical developmental period, concerning anatomical, neurochemical and behavioral changes. Moreover, adolescents are more sensitive to the long-term deleterious effects of drug abuse. Binge-like consumption of alcohol and marijuana, along with tobacco smoking, is a dangerous pattern often observed in adolescents during weekends. Nevertheless, the long-term effect of their adolescent co-exposure has not been experimentally investigated yet.

Long-Evans adolescent male (n = 20) and female (n = 20) rats from postnatal day 30 (P30) until P60 were daily treated with nicotine (0.3 mg/kg, i.p.), and, on two consecutive ‘binging days’ per week (for a total of eight times), received an intragastric ethanol solution (3 g/kg) and an intraperitoneal (i.p.) dose of cannabinoid 1/2 receptor agonist WIN55,212-2 (1.2 mg/kg). These rats were tested after treatment discontinuation at >P90 for associative food-rewarded operant learning in the two-lever conditioning chambers for six consecutive days on a fixed ratio 1 (FR1) schedule followed by another six days of daily FR2 schedule testing, after 45 days rest. We found the main effects of sex x treatment interactions in FR1 but not in FR2 experiments. Treated females show attenuated operant responses for food pellets during all FR1 and the FR2 schedule, whilst the treated males show an impairment in FR2 but not in the FR1 schedule. Moreover, the treated females’ percentage of learners was significantly lower than female controls in FR1 while treated males and females were lower than controls in FR2.

Our findings suggest that intermittent adolescent abuse of common drugs, such as alcohol and marijuana, and chronic tobacco exposure can cause significant long-term effects on motivation for natural reinforcers later in adulthood in both sexes. Females appear to be more sensitive to the deleterious effects of adolescent polydrug abuse with both sexes having an increased likelihood of developing lifelong brain alterations.

## Introduction

Adolescence is a critical period of brain development, concerning anatomical, neurochemical and behavioral changes. As the reward system has not yet matured, adolescents are particularly predisposed to risk- and novelties-seeking behavior that increases their likelihood of drug experimentation (Winters and Arria, 2011). Vulnerability to the effects of drugs of abuse during adolescence may be related to altered activity of the circuitry that mediates incentive processes and involves both dopamine (DA) and glutamate (GLU) (Burton et al., 2011). Compelling animal and human studies have shown that adolescent exposure to drugs, apart from acute damage (Sathanantham et al., 2021), may cause maladaptive changes in brain structures leading to the development of neuropsychiatric disorders such as anxiety, depression and substance use disorder in adult life (Salmanzadeh et al., 2020), that are different from those caused by adult drugs abuse.

The effects of drugs of abuse on the adolescent brain are so detrimental, that even if the exposure is limited to adolescence and then lifelong discontinued, it may still cause mental health disorders and alterations in adulthood supporting the theory of the presence of an age-dependent vulnerability of the brain to the drugs (Jordan and Andersen, 2017). Nevertheless, the long-term effects of drug abuse limited to the adolescent period have not constantly been observed in preclinical and clinical settings (Whyte et al., 2018).

Compelling evidence exists for adolescent binge drinking, i.e., consumption of a large quantity of alcohol (56-70 g) in a short period (≤2 h) (Abuse and Alcoholism, 2004), capable of inducing long-lasting alterations in brain development, plasticity and behavior (Lannoy et al., 2019), and increasing risks of psychiatric disorders i.e., alcohol use disorders in adult life (Crews et al., 2016).

Similarly, smoking (Leslie, 2020) or, now more commonly, vaping tobacco (Jenssen and Boykan, 2019) and cannabis (Jager and Ramsey, 2008; Vivian Chiu et al., 2022) in adolescent life seem to induce similar detrimental alterations in the maturing teen brain and with cannabis inducing disorders specifically in reward and motivation system (Pacheco-Colón et al., 2018).

An important problem when taking into account the long-term impact of adolescent drug addiction is that they are commonly co-abused and their deleterious effects may synergistically be potentiated (Pacheco-Colón et al., 2018; Singh, 2019). Indeed, adolescents often consume alcohol and cannabis in combination (Drugs and Crime, 2018) at weekends, while tobacco is commonly daily abused.

Nevertheless, the long-term effects of adolescent polydrug abuse have been less experimentally explored, with cannabis “binging” abuse never being investigated yet. Moreover, it is important to study if these impairments are sex-dependent, considering the well know gender effect in substance use disorder (Brady and Randall, 1999) and the recent attention to the inclusion of both sexes in animal experimentation (Voelkl et al., 2020). With this aim, we have set up a combined chronic (for nicotine) and binging (for alcohol and cannabinoid agonist) polydrug abuse administration protocol to study the specific impairments induced in reward and motivation in adult male and female Long-Evans rats. Specifically, adolescent (P30-P60) rats were treated with daily nicotine (0.3 mg/kg, i.p.), and, on two consecutive ‘binging days’ per week, with intragastric ethanol solution (3 g/kg) and the cannabinoid 1/2 receptor agonist WIN55,212-2 (1.2 mg/kg, i.p.) (Howlett et al., 2002). The rats were tested at >P90 for associative food-rewarded operant learning in the two-lever conditioning chambers for six consecutive days on a fixed ratio 1 (FR1) schedule followed by an interval of rest (45 days) and then by another six days of daily FR2 schedule to test eventual retention of reference memory impairment.

We found the main effects of sex x treatment interactions in FR1 but not in FR2 experiments. Treated females show attenuated operant responses for banana-flavored food pellets during all FR1 and the FR2 schedule, whilst the treated males show an impairment in FR2 but not in the FR1 schedule. Moreover, the treated females’ percentage of learners was significantly lower than female controls in FR1 while treated males were lower than male controls in FR2. However, the female-dependent susceptibility to adolescent drug abuse can also be due to an impairment of the DAergic and reward system that involves dysregulation of other areas such as BLA and mPFC/CCG (Walker et al., 2017). Our findings suggest that adolescent polyabuse of common drugs such as tobacco and binge-like intoxication of alcohol and marijuana can cause significant long-term effects on motivation for natural reinforcers later in adulthood, especially in females.

## Methods

### Animals

Twenty male and twenty female Long-Evans rats (about 28 day-olds) were obtained from a colony bred at the University of Malta. The drug treatment started a P30 and ended at P60 while they were tested at age >P90. Animals were housed in a 12:12 light cycle (lights on at 07.00 a.m. and off at 07.00 p.m.). Purified tap water and food chow were available ad libitum throughout the course of the study, except when the animals were exposed to food deprivation. All animal procedures were carried out under the University of Malta’s ethical guidelines and in conformity with Maltese and international laws and policies (EU Directive, 2010/63/EU for animal experiments). All efforts were made to minimize animal suffering and to reduce the number of animals used.

### Pharmacological treatment and experimental design

#### Drugs

Ethanol 95% (v/v) and (-)-nicotine hydrogen tartrate salt ((-)-1 -Methyl-2-(3-pyridyl) pyrrolidine (+)-bitartrate salt were purchesed by Sigma Aldrich (St. Louis, MO, USA). The CB1/2 receptor agonist (R)-(+)-[2,3-Dihydro-5-methyl-3-(4-morpholinylmethyl)pyrrolo[1,2,3-de]-1,4-benzoxazin-6-yl]-1-naphthalenylmethanone mesylate, (R)-(+)-WIN 55,212-2) WIN 55,212-2 was purchased from Tocris Cookson Ltd. (Bristol, United Kingdom). Ethanol was intragastric (i.g.) administrated as a 25% (v/v) ethanol solution in water. WIN 55,212-2 was freshly dissolved in a vehicle solution (2□ml/kg) made of 5% PEG-400, 5% Tween 80 in saline and i.p. administered. Nicotine was dissolved in saline, with pH adjusted to about 7.4. and i.p. administered and the weight was given as free-base. In our study, we used the intermediate 0.33 mg/kg nicotine dose capable to enhance reward function and psychomotor performance in adolescent and adult male and female rats (Xue et al., 2020) and induce anxiety (Casarrubea et al., 2015; Casarrubea et al., 2021; Casarrubea et al., 2020) and dopamine cell excitation in adult rats (Pierucci et al., 2022). The dose of 3 g/kg alcohol, i.g., administered, was chosen because it is comparable to previously reported data (Helfer et al., 2009; Lauing et al., 2008; Maldonado-Devincci et al., 2010), and capable of inducing long-term alterations in memory and behavior (Mooney-Leber and Gould, 2018). Moreover, we used a dose of WIN55,212-2 (1.2 mg/kg), higher than the dose self-administered by rats (Fattore et al., 2007; Kirschmann et al., 2017), that was capable to induce deficits in short-term memory when chronically administered in adolescent male rats (O’Shea et al., 2006; Schneider and Koch, 2003) but not in female rats (Kirschmann et al., 2017) and impaired the dentate gyrus LTP in adult male rats (Colangeli et al., 2017). The dose of WIN55,212-2 used here can be considered equivalent to a cannabis cigarette (~800 mg) containing a dose of 3 mg/kg THC (~20% Δ9-THC) (Health Canada, 2013), considering that the synthetic cannabinoid agonist has a higher affinity for CBRs than THC (Lawston et al., 2000).

#### Experimental Protocol

The experimental protocol was designed to mimic the poly-abuse pattern typical of teenagers/young adults (Figure 1A). The binge-drinking alcohol (3 g/kg, intragastric i.g.) with the contextual WIN 55,212-2 (1.2 mg/kg, i.p.) administration and daily nicotine administration (0.3 mg/kg, i.p.) mimicked the 2-day heavy drinking and moderate abuse of cannabis in light smoking teenagers at the weekend. Two groups of male and female adolescent rats (n = 40), from P30 to P60, were weekly exposed to a two-day binge of administration of 25% (v/v) ethanol solution or its vehicle for four consecutive weeks. On the same days, rats received an i.p. low/moderate dose of WIN 55,212,2. Moreover, for the period of observation (30 days) the rats received a daily low dose of nicotine administration (0.3 mg/kg, i.p.) (Matta et al., 2007) (Figure 1A). To reduce the stress of the manipulations, rats were handled for a week before the experimentation.

**Figure 1:**
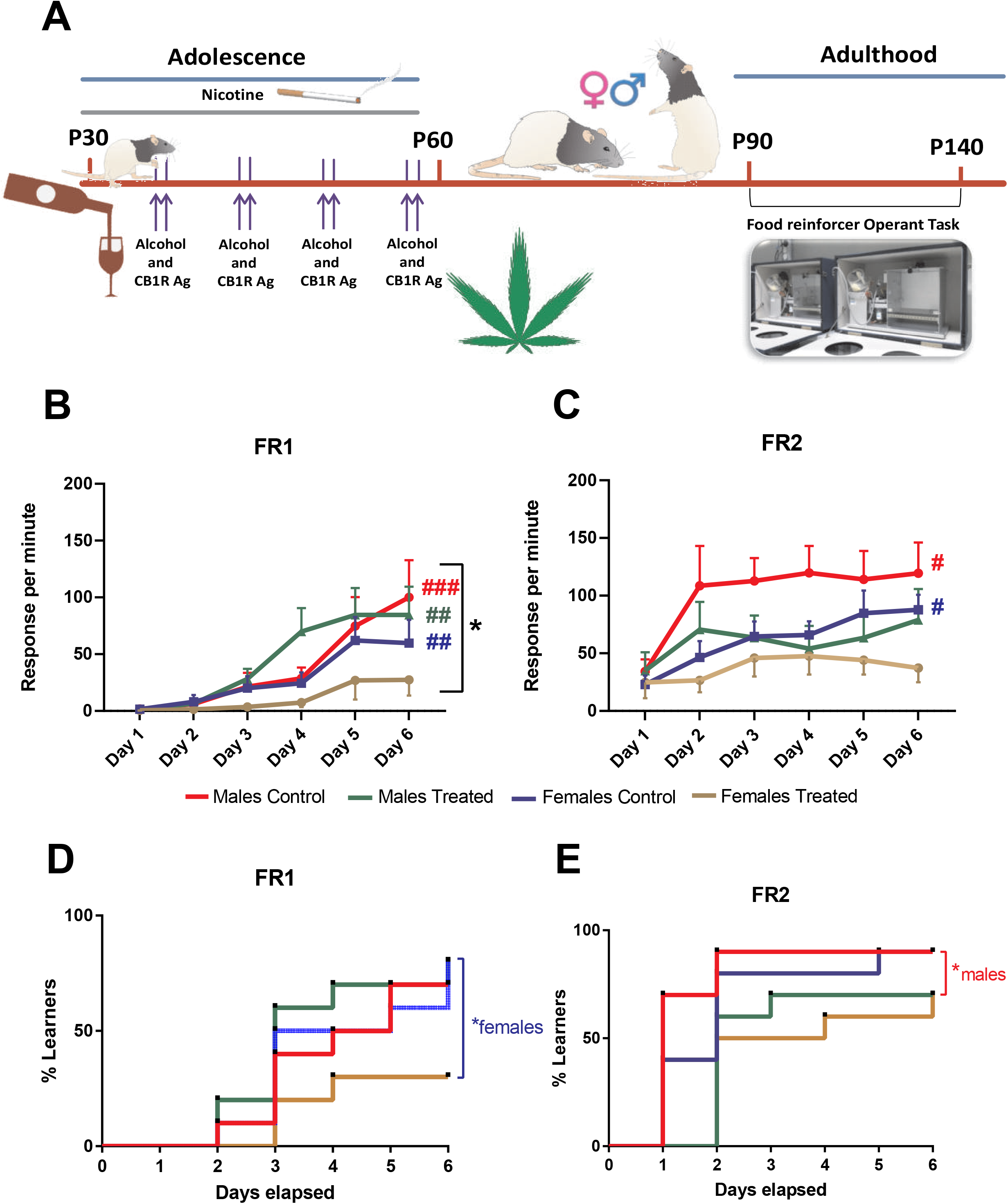
Long-term term effect of adolescent binging on alcohol and cannabinoid and daily exposure to nicotine on learning and motivation in rats. **A.** The chronic treatment was initiated at P30 with daily nicotine administration (0.3 mg/kg., i.p.) for 30 days and two consecutive days once a week with ethanol (3 g/kg; i.g.) and 1.2 mg/kg CB 1/2 receptor agonist WIN 55,212-2. Rats were food restricted before each test until they reached 85% of their ad libitum body weight. Operant conditioning testing was performed at least after 30 days the end of the treatment (>P90). **B.** Rats underwent the first 6 days of testing under a FR1 schedule. Female rats made fewer presses compared to males. Control males, treated males and control females improved throughout the FR1 schedule when compared to FR1 day 1, while treated females did not. **C.** After 45 days, rats underwent to a FR2 schedule. Male and Female controls improved in response per min during the FR2 while both treated male and female groups did not show an improvement during the six days of the test. *p<0.05, Two-way ANOVA; #p<0.01; ##p<0.005; ###p<0.001generalized Kruskal Wallis analysis. **D.** Instrumental learning performance was significantly decreased in poly-treated females during FR1 and in (**E.**) poly-treated males during FR2 compared to their respective control groups. *p<0.05, log-rank Mantel-Cox test. i.p.: intraperitoneal; i.g.: intragastric; FR: fixed ratio.

#### Operant Instrumental Learning

Eight operant conditioning chambers (Lafayette Instrument, Lafayette, IN) were used, in a dedicated temperature and humidity control laboratory. In all operant conditioning sessions, one lever was designated as the active lever and remained active during all sessions and the second lever was designated as inactive and left retracted. The food reinforcer was a 45 mg appetitive pellet (Precision Sucrose pellet, Banana flavor-LBS LTD). Rats were tested at age >P90 and submitted to a one-time random feeding as a training session. All training sessions were performed one day prior to the FR1 test. These consisted of 25 randomly timed pellets that were dispensed from the roulette through the tube and into the feeder without the need for the rat to press the lever. The time, which was randomly chosen by the program, was not less than 2 seconds but no more than 60 sec, and with a 20 sec delay time. Rats were food restricted before each test until they reached 85% of their ad libitum body weight (Zellner and Ranaldi, 2006). The food restriction was maintained throughout the testing and the interval between FR1 and FR2. During the food deprivation period, the rats were handled daily for about 10 min. Rats were left in their cage for about ten min and placed in the operant conditioning chambers for another ten min as a habituation procedure before being tested. Rats underwent the first 6 days of testing under an FR1 schedule. The rats had 60-sec chance to press the level and if successful a food pellet was provided as a reward. A 20-sec delay followed each trial with the stimulus light on. This was repeated 25 times. After a resting period, with their body weight to 85% of their ad libitum weight, the rats were tested for another 6 days of FR2 schedules. The same parameters were used as in the FR1 but with the need for two presses for one reward. The data collected were the responses of the rats on the lever to obtain food per 25 schedules and also the latency in seconds for the pressing of the lever. If the lever was not pressed in the permitted time interval, this was taken as a full latency of 60 seconds. For data processing, the response per min or rate of response was calculated from the two parameters obtained from the test by using the equation below (Beninger, 1991).

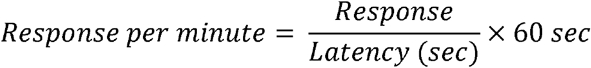

Differences between learners and non-learners between treatment groups of both female and male groups were also calculated. All the treatment groups during all test days were compared to identify if the treatment created any difference in associative learning. The analysis of this data also produced a distinction between the days of the test, and hence a measure of the progress achieved by rats. Learners were classified as such when they achieved a 100% response within the test schedule. One of the dependent variables was the learning outcome (Learners vs non-learners) while the independent variables were the day of the test, the sex and the treatment regime.

#### Statistical analysis of the data

ABET II software (Lafayette Instrument, Lafayette, IN) was used to control, monitor and record individually from all chambers via an independent interface.

A three-way ANOVA analysis was performed so that an interaction effect between the three independent variables on the continuous dependent variable could be determined if it exists. So here it was important to identify if there was an interacting effect of the drugs under study on the associative learning potential of the rats, together with sex and day of the test. So a possible three-way interaction effect could be determined between the three independent variables. A statistically significant three-way interaction was determined by a p < 0.05 criterion. A two-way ANOVA analysis followed, to compare the mean differences between groups (sex x day of the test), (sex x treatment) and (day of test x treatment) so that it could be understood if there was an interaction between all combinations of the two independent variables on the associative learning potential in rats.

Data that did not display equal variances were analyzed using nonparametric tests. Kruscal-Wallis non-parametric statistical test was performed to test the significant difference between the responses per min output of each day when compared with its starting point, for each sex under the two different treatment conditions.

The performance of each rat to reach the learning criterion was analyzed using Kaplan-Meier event analysis over the instrumental learning period, and the resulting curves were compared by employing the log-rank Mantel-Cox test. Statistical analysis was carried out using GraphPad Prism v. 9 (GraphPad Software, Inc., San Diego, CA). All values represent the mean ± standard error of the mean. A P-value lower than 0.05 level was considered significant.

## Results

### Effect of the adolescent polydrug treatment on weight change

All rats were housed in the same room with good and adequate environmental conditions with food and water ad libitum. The weight of each individual rat was recorded regularly from P30 to P90. The weight gain was smaller for the treated group compared to the control group for both males (−54.5 ± 13.36 g; F(18)=0.715, p=0.001) and females (−22.8 ± 9.56 g; F(18)=0.166, p=0.028; not shown). No difference instead was observed between the weight gain between males and females (p>0.5; not shown).

### Long-term effect of the adolescent polydrug treatment on operant food-conditioning acquisition

The effects of adolescent exposure to our polydrug treatment on the adults’ instrumental learning were assessed by evaluating the animal’s ability to acquire simple instrumental tasks FR1 and, after 45 days interval, FR2.

First, we compared the performance of the 4 groups considering the response, that it is equal to the active lever presses per minute Three-way ANOVA analysis did not show significant three-way interaction between sex, day of test and treatment for the associative learning potential during FR1 (F(5, 216)=0.569, p>0.05). A significant two-way interaction was found for sex x treatment (F(1, 216)=4.384, p<0.05). No significant two-way interaction was found for the day of test x treatment (F(5, 216)=0.671, p>0.05) and for sex x day of the test (F(5, 216)=1.819, p>0.05) (Fig. **1B**). No statistically significant three-way interaction between Sex, day of test and treatment was found for the associative learning potential during FR2 (F(5, 216)=0.300, p>0.05). No significant two-way interaction was found for sex x treatment (F(1, 216)=1.111, p>0.05). No significant two-way interaction was found on the day of test x treatment (F(5, 216)=0.897, p>0.05). No significant two-way interaction was found for sex x day of the test (F(5, 216)=0.553, p>0.05).

When considering the learning performance, generalized Kruskal Wallis analysis showed that control males improve throughout the FR1 schedule when compared to FR1 day 1 (χ2(11)=43.980,p<0.001). This improvement was also seen during the whole FR2 schedule (χ2(5)=11.778,p=0.038). Treated males showed an improvement during the FR1 schedule (χ2(5)=15.658,p=0.008) Of note, treated males showed no improvement during the FR2 schedule (χ2(5)=1.698,p=0.889). Control females showed a generalised improvement during FR1 schedule (χ2(5)=17.614,p=0.003) and FR2 schedule (χ2(5)=12.984,p=0.024). Treated females showed no improvement in response per min during the FR1 schedule (χ2(5)=0.545,p=0.990) and the FR2 schedule (χ2(5)=2.093,p=0.836).

Remarkably, the memory retention (expressed as the difference in active lever responses between FR1 day 6 and FR2 day 1) after the interval period (7 weeks) between the two schedules was similar for all the groups, except the treated females who showed a very similar value (Treated female FR1 day six 27.39 ± 13.78 and 24.79 ± 13.71 (χ2(1)=0.073,p=0.787). This was because treated female rats ended at a very low level of associative learning on the 6^th^ day of FR1. Therefore, the present findings indicate that the acquisition of the present task was soo impaired in female-treated rats that no retention was possible.

Moreover, we analysed the instrumental learning performance in terms of days elapsed to reach the learning criterion for each subject within the 4 groups for both FR1 and FR2 schedules. Kaplan-Meier analysis showed for FR1 that median learning times were three days for all the groups apart from female poly-treated rats. Indeed, 30% of male and female control rats and 20% of males and 70% of females of the poly-treated rats failed to reach the criterion within 6 days. The log-rank Mantel-Cox test for comparison of survival curves indicated that FR1 learning performance was significantly decreased in poly-treated females compared to their control rats (χ2 = 4.39, df = 1, p = 0.0360; figure 1D). Kaplan-Meier analysis for FR2 showed that median learning times were one day and two days for male and female controls, respectively, two days and one day for poly-treated males and females, respectively. In addition, 10% of male and female control rats and 30% of males and females of the poly-treated rats failed to reach the criterion within 6 days. The log-rank Mantel-Cox test for comparison of survival curves indicated that FR2 learning performance was significantly decreased only in poly-treated males compared to control rats (χ2 = 6.12, df = 1, p = 0.0134; figure 1E).

## Discussion

Nicotine, alcohol and cannabis are the most commonly abused substance among youth (World Drug Report, 2022), most likely due to their easy obtainment. Although these substances are generally co-abused by adolescents, little is known about their combined longterm effects on the brain and health in general. Indeed, most of the studies have taken into consideration the short- and long-term effects of a single drug administered, prevalently chronically, during adolescence (see for a recent review (Mooney-Leber and Gould, 2018)). Moreover, alcohol and cannabis are frequently used by teenagers and young adults in a binge pattern, particularly during the weekend (Drugs and Crime, 2018). While the short and longterm effect of binge drinking during adolescence has been intensively investigated on brain functions (Pérez-García et al., 2022), the consequences of cannabis binges, typical of social situations when there is the ready availability of cannabis, remain unknown.

The first outcome of this study is about the short and long-term effects of the polydrug treatment on the body weight of the rats. It is well known that drug abuse affects food intake and body weight (Mohs et al., 1990), but conflicting results exist instead about the effect of their adolescent administration. For instance, adolescent nicotine exposure does not affect adolescent and adult weight gain (Pushkin et al., 2019) but reduced the nicotine anorectic effect (Natividad et al., 2013) in adulthood. On the other hand, binging on 3 g/kg alcohol for 4 weeks produced a reduction (Lauing et al., 2008) or no change (Helfer et al., 2009) in the post-treatment body weight in rats. Conflicting results with lack of effect or decrease of body weight have been obtained with adolescent cannabis smoke exposure from P35-P45 ((Bruijnzeel et al., 2019; Hernandez et al., 2021), or to WIN55,212-2 (P30-P43) (Pushkin et al., 2019; Schoch et al., 2018). Conversely, in the current study, the rat body weight gain during the polydrug treatment at P30-P60 but also a P90 was, as expected, higher in males compared to females but independently from the adolescent exposure to the drugs.

The lack of the anorectic effect of our polydrug treatment on body weight gain is probably due to either the particular binging paradigm of administration used here and/or the counteracting effects of the single drugs. In support of the latter hypotheses, a slightly higher dose of WIN55,212-2 (2 mg/kg) produced a decrease in body weight gain, although only in female adolescent mice, that was prevented by the co-exposure with 0.36 mg/kg nicotine (Pushkin et al., 2019).

As far as our findings about associative food-rewarded operant learning are concerned, they are in line with a large body of evidence showing a sex-dependent alteration of the brain reward system (Walker et al., 2017) and related learning, induced by adolescent exposure to drugs of abuse, with the females being the most affected (Mooney-Leber and Gould, 2018). Indeed, we found a decreased effect of natural rewards (i.e., the palatable banana-flavored pellets) on inducing instrumental learning that was sex-dependent although both sexes were differently affected. After a period of 30-day washout from the last polydrug treatment (at >P90), over the FR1 schedule of reinforcement, only treated females were not able to acquire and maintain the instrumental conditioning task (i.e., lever pressing). It is interesting to note, there was an expected decrease in level-pressing values of FR2 Day 1 compared to FR1 Day 6 due to the resting time between the two tests, an expression of memory retention that facilitates re-acquisition of a similar already acquired task (see (Ohta et al., 1993)), which resulted similar for all the groups apart from the females showing no change. Indeed, the polydrug treatment completely hindered female rats’ acquisition of the task acting on the brain circuitry sustaining instrumental learning (Balleine and Dickinson, 1998). This includes the dopamine (DA)-containing neurons of the ventral tegmental area (VTA) projecting to the core of the nucleus accumbens (NAc) (Chase et al., 2015; Heymann et al., 2020) and other areas receiving DAergic innervation i.e., dorsal medial striatum (Yin et al., 2005), amygdala (Andrzejewski et al., 2005), mPFC and anterior cingulate cortex (ACC) (Baldwin et al., 2002; Caballero et al., 2019). Moreover, another neurotransmitter system impacted by the polydrug treatment is glutamate (GLU) and its NMDA receptors which have a clear role in the acquisition of instrumental learning (Freed and Wyatt, 1981) interacting with DA as, for instance, the genetic deletion of NMDA receptors in VTA DA neurons slowed instrumental learning (James et al., 2015).

Indeed, adolescent exposure to chronic alcohol (Sircar and Sircar, 2006) or binge drinking (Obray et al., 2022) and Δ9 THC (Rubino et al., 2015) depress D1 and NMDA receptor signals in adults. A reduction of the DA D1 and NMDA receptor function in the NAc in females compared to males could explain our findings in animals prior exposed to polydrugs. Consistently, D1 receptor antagonist SCH23390 (Hernandez et al., 2005; Smith-Roe and Kelley, 2000) and NMDA receptors antagonist AP-5 (Kelley et al., 1997) infusion in the NAc core impaired both instrumental learning and performance but did not affect memory consolidation and retrieval (Hernandez et al., 2005). Similar results were obtained with the blockade of NMDA receptors by AP-5 infusion in the basolateral amygdala (BLA), mPFC (Baldwin et al., 2000) and ACC (McKee et al., 2010) completely abolished acquisition while the retention of lever pressing for food response was unaffected. A rise of accumbal DA release occurs during the initial acquisition of the task (Segovia et al., 2011), proportionally to the rates of lever-pressing (Salamone et al., 1994). The alteration of the acquisition of FR1 and the maintenance of the instrumental response in female rats might depend on the lack of selective increase in the firing activity of VTA neurons projecting to the core NAc (Heymann et al., 2020). Consistently, adolescent Δ9 THC (Scherma et al., 2016) WIN55,212-2 (Pistis et al., 2004) exposure decreased the VTA DA neurons’ firing activation and NAc DA release induced by WIN55,212-2 administration in adult rats (Scherma et al., 2016) and induced a functional tolerance of VTA DA neurons to WIN55,212-2, morphine, amphetamine and cocaine administration on the DA neuron firing rate (Pistis et al., 2004).

Late adolescent (P40-P65) 1.2 mg/kg WIN55,212-2 treatment decreased palatable food intake and breakpoint under progressive ratio (PR) in adulthood after a wash-out of 20 days (Schneider and Koch, 2003). Oppositely, early adolescent (P30-43) exposure to WIN55,212-2 (Schoch et al., 2018) or cannabis smoke (Hernandez et al., 2021) did not affect food motivation under a PR schedule of instrumental responding for food reward at >P70. As far as nicotine, a similar dose (0.35 mg/kg) to that used in the current study daily administered to adolescent rats (P35-50) was found to induce better reward-related learning in adult males while impairment in female rats (Quick et al., 2014), but resulted ineffective when administered in adult mice (Pushkin et al., 2019). Finally, alcohol abuse during adolescence is known to cause impairment in learning and memory in adult life in rats (White et al., 2000) and humans (Crews et al., 2019; Crews et al., 2016) but initial instrumental learning is generally unaffected (Risher et al., 2013).

The only available evidence of adolescent drug abuse co-exposure on instrumental learning showed that prior 0.36 mg/kg nicotine and 2 mg/kg WIN55,212-2 treatment did not change the ability of both sexes to learn an operant task to obtain food reward in adult mice (Pushkin et al., 2019). Our polydrug treatment also included binge drinking, and this might be responsible for the diverse outcome.

Our findings indicate that control male and female rats showed good performance when retested in the FR2 schedule, while both treated males and females did not improve their performance of the represented task. Consistently, the percentage of the learners (those who meet the criterion) observed in the control males in FR2 was higher than in FR1, reaching the maximum of the learners on Day 2 FR2.

Our findings would indicate that female-treated rats showed deficits in the acquisition, performance (FR1), and consequently in the consolidation of the reference memory (FR2), while treated males have a selective deficit of re-acquisition and performance in FR2 with a normal memory process in FR1.

The reason for this selectively delayed male deficit on FR2 induced by the polydrug treatment compared to females is not evident. It might be related to the different sex-dependent neuronal reorganization and DA and GLU receptor pruning that occur during adolescence (Walker et al., 2017) or aging (FR2 occurred after 45 days) (Ohta et al., 1993) and/or a specific learning deficit upon retrieval in the reconsolidation (Milekic and Alberini, 2002). Other neurotransmitter systems cannot be rouled out, for instance serotonin (5-HT) for its implication in memory and reward (Bombardi et al., 2021b; De Deurwaerdere and Di Giovanni, 2020) and its interaction with nicotine (Bombardi et al., 2021a; Bombardi et al., 2020; Bombardi and Di Giovanni, 2013; Venzi et al., 2016). These possibilities urge experimental validation.

In conclusion, this is the first evidence showing that smoking and having a few drinks and a joint at the weekend, which is considered not particularly harmful by adolescents/young adults, has instead long-term gender-dependent effects on the learning and reward system. The extrapolation of our data to humans further supports the concept of adolescence as an ideal time to employ gender-targeted preventive and meliorative (or primary and secondary preventive) strategies, instead of relying on tertiary prevention in adulthood. Considering that adolescence has been referred to as the gateway to adult health outcomes (Raphael, 2013) it would be pivotal to act fast on prevention.

## AUTHORS CONTRIBUTION

Conceptualization, G.D.G.; methodology, N.A. and G.D.G.; formal analysis, N.A. and G.D.G.; investigation, K. H. and N.A.; resources, G.D.G.; data curation, N.A. and G.D.G.; writing—original draft preparation, writing—review and editing, G.D.G., K. H. and N.A.; supervision, G.D.G.; project administration, G.D.G.; funding acquisition, G.D.G. All authors have read and agreed to the published version of the manuscript.

## CONFLICT OF INTEREST

The authors declare that the research work was conducted in the absence of any commercial or financial relationships that could be construed as a potential conflict of interest.

## ACKNOWLEDGMENTS

Our research was supported by the University of Malta, Faculty of Medicine Research Support Grant (N. A.). and an Erasmus fellowship (K. H.). The assistance provided by Dr Massimo Pierucci and Ms Maria Vella was greatly appreciated.

## Notes

### Competing Interest Statement

The authors have declared no competing interest.

